# The role of viscoelasticity in long-time cell rearrangement

**DOI:** 10.1101/2021.08.09.455610

**Authors:** Ivana Pajic-Lijakovic, Milan Milivojevic

## Abstract

Although collective cell migration (CCM) is a highly coordinated and ordered migratory mode, perturbations in the form of mechanical waves appear even in 2D. These perturbations caused by the viscoelastic nature of cell rearrangement are involved in various biological processes, such as embryogenesis, wound healing and cancer invasion. The mechanical waves, as a product of the active turbulence occurred at low Reynolds number, represent an oscillatory change in cell velocity and the relevant rheological parameters. The velocity oscillations, in the form of forward and backward flows, are driven by: viscoelastic force, surface tension force, and traction force. The viscoelastic force represents a consequence of inhomogeneous distribution of cell residual stress accumulated during CCM. This cause-consequence relation is considered on a model system such as the cell monolayer free expansion. The collision of forward and backward flows causes an increase in cell packing density which has a feedback impact on the tissue viscoelasticity and on that base influences the tissue stiffness. The evidence of how the tissue stiffness is changed near the cell jamming is conflicting. To fill this gap, we discussed the density driven change in the tissue viscoelasticity by accounting for the cell pseudo-phase transition from active (contractile) to passive (non-contractile) state appeared near cell jamming in the rheological modeling consideration.

## 1. Introduction

Collective cell migration (CCM) influences the viscoelasticity of multicellular systems which has a feedback impact to a long-time cell rearrangement and the biological and mechanical state of single cells. Deeper insight into this inter-relation by emphasizing the role of viscoelasticity is essential for understanding of various biomedical processes such as wound healing, morphogenesis, and tumorigenesis (Blanchard et al., 2019; Barriga et al., 2018; Barriga and Mayor, 2019; Pajic-Lijakovic and Milivojevic, 2019a,b). The role of viscoelasticity in a long-time cell rearrangement will be discussed on the model system such as a cell monolayer free expansion. The CCM induces a generation of the mechanical waves (Serra-Picamal et al., 2012; Tlili et al., 2018; Petrolli et al., 2021). The mechanical waves represent oscillations of cell velocity and the rheological parameters of multicellular system. The generation of mechanical waves is closely related to a viscoelasticity at supracellular level (Pajic-Lijakovic and Milivojevic, 2020c). The phenomenon represents a part of elastic turbulence occurred during the flow of soft matter systems under low Reynolds number. Grosman and Steinberg (1998;2000) considered the elastic turbulence appear during the externally induced flow of flexible, long-chain polymer solutions by introducing the Weissenberg number 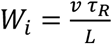 as relevant (where is the velocity, is the characteristic length, and is the stress relaxation time). In contrast to other soft matter systems, the multicellular systems are active, capable of inducing the self-rearrangement which has been treated as an active turbulence (Alert et al., 2019). Cell velocity oscillations during the monolayer free expansion appear in the form of local forward and backward flows (Serra-Picamal et al., 2012). While the forward flow is caused by chemotaxis, haptotaxis, durotaxis (Murray et al., 1988), the backward flow is driven by the viscoelastic force and the surface tension force (Pajic-Lijakovic and Milivojevic, 2020c). The viscoelastic force represents a consequence of the inhomogeneous distribution of cell residual stress accumulated during CCM. It is a resistive force acts always opposite to the direction of cell movement. The surface tension force acts to reduce the cell monolayer surface.

The viscoelasticity caused by CCM has been considered on two time scales: (1) a short-times cale (a time scale of minutes) and (2) a long-time scale (a time scale of hours) (Pajic-Lijakovic and Milivojevic, 2019a;2020b,c). CCM occurs on a long-time scale and induces the generation of strains (volumetric and shear), as well as their long-time change. These strains cause the generation of corresponding stresses (normal and shear), their short-time relaxation, and long-time residual stresses accumulation.

Collision of forward and backward flows can induce a local increase in cell packing density and on that base change a viscoelasticity and even induces the cell jamming state transition. Nnetu et al. (2012) and Tlily et al. (2018) pointed to fluctuations of cell packing density during the cell monolayer free expansion. Pajic-Lijakovic and Milivojevic (2021b) and Pajic-Lijakovic (2021c) discussed density driven changes in the viscoelasticity of multicellular systems. They discussed that the density-driven evolution of viscoelasticity induces several transitions between five viscoelastic states gained within three regimes: (1) convective regime, (2) conductive regime, and (3) damped-conductive regime (Pajic-Lijakovic and Milivojevic, 2021b). The convective regime accounts for two states of viscoelasticity depending on the magnitude of cell velocity and the state of cell-cell adhesion contacts. Higher cell velocities, i.e.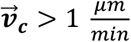 and weak cell-cell adhesion contacts (characteristic for lower cell packing densities) lead to pronounced liquid-like behavior described by the Maxwell model, while lower cell velocities 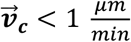and stronger cell-cell adhesion contacts (characteristic for confluent multicellular systems) ensure solid-like behavior described by the Zener model. The Maxwell model was chosen based on experimental data of long-time viscoelasticity of cell aggregate under micropipette aspiration reported by Guevorkian et al. (2011). The Zener model was chosen based on experimental data of (1) the MDCK cell monolayer free expansion (Serra-Picamal et al., 2012) and (2) CCM of the confluent MDCK cell monolayer (Notbohm et al., 2016). This model also corresponds to long-time viscoelasticity of cell aggregate uni-axial compression between parallel plates considered by Marmottant et al. (2009) and Mombash et al. (2005). Transition from convective to the conductive regime is accompanied with significant reduction in cell mobility caused by increasing the normal residual stress accumulation. This so called “plateau” regime near the system jamming described by the Kelvin-Voigt model corresponds to the velocity 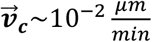 (Pajic-Lijakovic and Milivojevic, 2017;2021b). Further increase in cell packing density leads to an anomalous nature of energy dissipation during CCM as a characteristic of the damped-conductive regime (Nnetu et al., 2013). This regime accounts for two sub-regimes of viscoelasticity, i.e. the cell transient sub-regime for the cell velocity 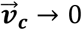 and the jamming sub-regime for the cell velocity 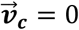 described by the fractional constitutive model (Pajic-Lijakovic and Milivojevic, 2019c;2021b). Braumgarten and Tighe (2017) reported that systems have to pass through a transient regime before reaching the jamming state. We expanded our previous consideration (Pajic-Lijakovic and Milivojevic, 2021b) by accounting for cell pseudo-phase transition from active (contractile) to passive (non-contractile) state in the viscoelasticity of the transient sub-regime.

The density driven changes in the viscoelasticity induce the system stiffening. The stiffness increases with cell packing density if and only if cells keep their active (contractile) state. However, near jamming cells undergo the pseudo-phase transition from active (migrating) to passive (resting) cell state under constant cell packing density (Bi et al., 2015;2016; Pajic-Lijakovic and Milivojevic, 2019c). Despite extensive research devoted to the study the cell jamming state transitions, we still do not fully understand the phenomenon from the standpoint of the tissue stiffness change. This contribution represents an attempt to clarify this issue. This pseudo-phase transition influence the mechanical state of single cells (Pajic-Lijakovic and Milivojevic, 2019a;2020a,b). Zimmermann et al. (2016) considered the cell jamming state transition during CCM of MDCK cell monolayers and revealed that cells under jamming state actively down-regulate their propulsion forces in response to an increase in cell packing density by discussing the contact inhibition of locomotion (CIL) as the main mechanisms. Garcia et al. (2015) indicated several effects recognized within a cellular system near jamming: (1) a decrease in the propulsive force produced by each cell, (2) an increase in the friction of the cells with the substrate, and (3) increase in the cell−cell effective friction. Bi et al. (2015) quantified the cell jamming state transition by the cell shape parameter. Near jamming, cells change the shape factor from the elongated shape characteristic for motile cells to the less elongated characteristic for passive cells.

Active (contractile) cells are much stiffer than passive (non-contractile) ones due to an accumulation of contractile energy (Kollmannsberger et al., 2011; Pajic-Lijakovic and Milivojevic, 2019a; Schulze et al., 2017). Schulze et al. (2017) revealed that Young’s modulus of contractile MDSC cell monolayer is ∼33.0 ± 3.0 *kPa*, while the modulus of non-contractile cells is significantly lower and equal to ∼15.6 ± 5.5 *kPa*. However, many authors neglected single-cell softening during the pseudo-phase transition near the cell jamming such as Bi et al. (2015;2016), Merkel and Manning (2018) and Lawson-Keister and Manning (2021) and described the cell jamming state as the stiffest cell state. Under the jamming state multicellular system reaches a maximum cell packing density, but the cells themselves become passive (non-contractile) and on that base softer. The main goal of this contribution is to clarify this issue by (1) discussing the impact of the density driven change in the viscoelasticity on the stiffness of the multicellular systems and (2) accounting for pseudo-phase transition into viscoelasticity for cells near jamming.

## 2. Free expansion of cell monolayer: the generation of forward and backward flows

The cell long-time rearrangement during a monolayer free expansion is guided by several forces such as (1) the viscoelastic force, (2) the surface tension force, and (3) the traction force (Murray et al., 1988; Pajic-Lijakovic and Milivojevic, 2020c). The viscoelastic force represents a consequence of an inhomogeneous distribution of cell residual stress on one hand and matrix residual stress on the other, and is equal to 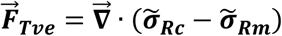(where 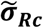 is the cell residual stress and 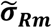 is the matrix residual stress) (Murray et al., 1988; Pajic-Lijakovic and Milivojevic, 2020c). The viscoelastic force is the resistive force and acts always opposite to the direction of cell movement in order to suppress it. The surface tension force is equal to 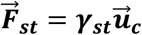 (where ***γ***_*st*_ is the surface tension and 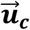 is the local cell displacement field). This force acts to reduce the surface of cell monolayer. The traction force is equal to 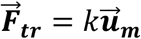 (where *k* is an elastic constant of cell-matrix adhesion contact and 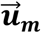 is the matrix displacement field exerted by cells) and acts in the direction of cell movement. This force influences the rate of cell expansion depending on the rheological behavior of a matrix (Murray et al., 1988; Pathak et al., 2012; Notbohm et al., 2016). Higher traction force is generated at a stiffer matrix (Pathak et al., 2012). Cells undergo free expansion in the form of forward flows performed in two opposite directions, i.e. and *x* ∈ (0, *L*(*τ*) (where *L*(*τ*) is the half of the monolayer length) (Figure 1).

**Figure 1.**
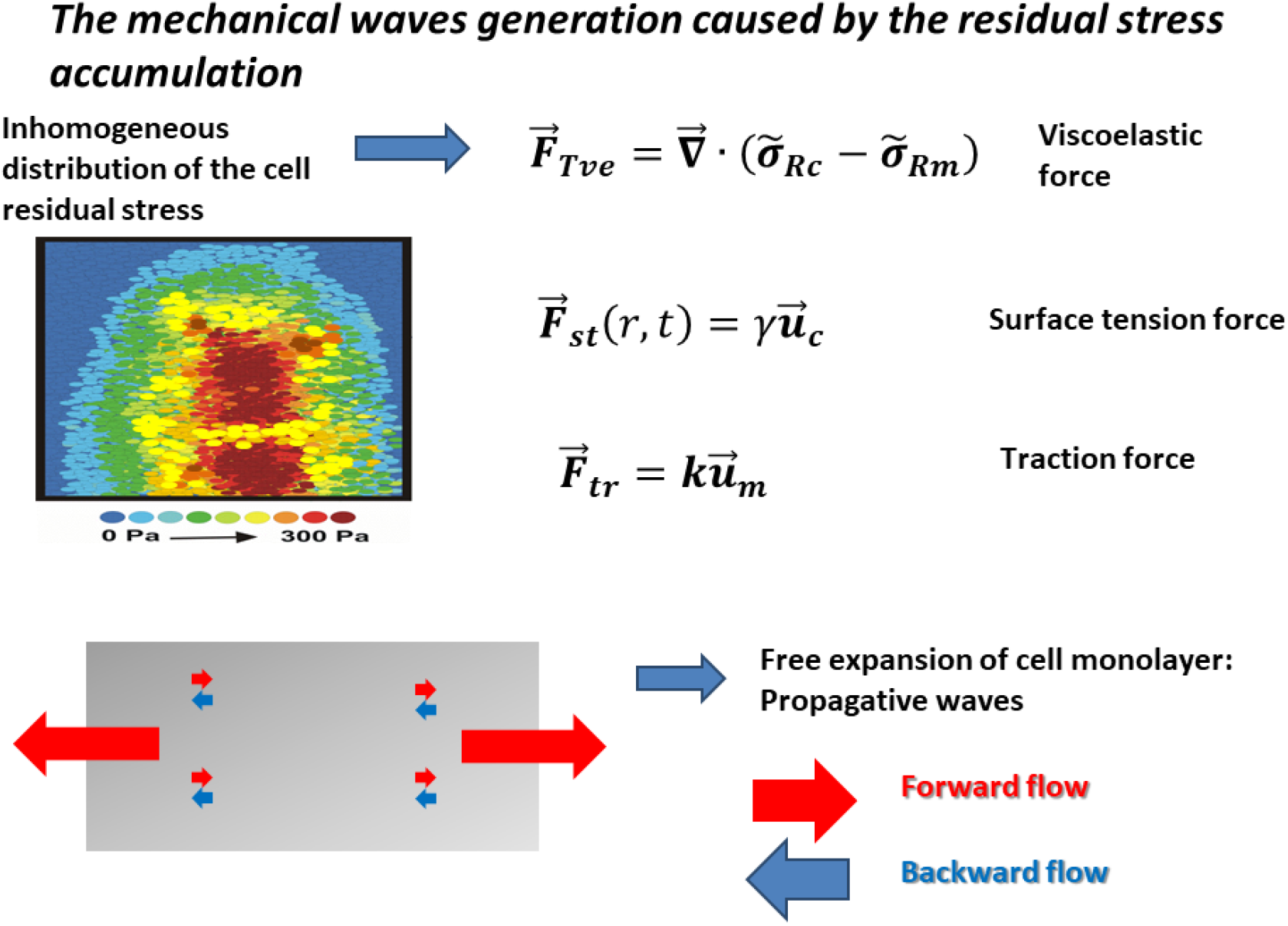
The schematic presentation of the forward and backward flows generated during the cell monolayer free expansion.

The forward flow driven by chemotaxis (for the cell velocity 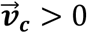) leads to an increase in: the viscoelastic force and the surface tension force capable of reducing further cell movement. An increase in the viscoelastic force, as a consequence of the cell residual stress accumulation, causes the monolayer local stiffening. When these two forces reach maximum values, the cell velocity is 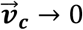. This condition corresponds to a generation of the backward flow (for the cell velocity 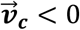) driven by the viscoelastic force and the surface tension force. The backward flow leads to a local softening of the monolayer and corresponding decrease in both forces. A decrease in these forces reduces the backward flow. Then, the forward flow starts again. Simultaneous generation of forward and backward flows induces oscillatory changes in cell velocity and has been discussed in the context of apparent inertial effects (Alert et al., 2020; Pajic-Lijakovic and Milivojevic 2020c). The schematic presentation of the forward and backward flows generated during the cell monolayer free expansion is given in Figure 1.

The cell velocity represents the rate of change the cell displacement field 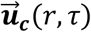 equal to:

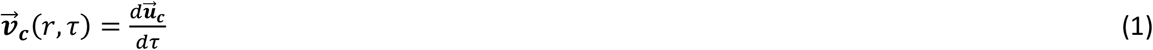

where *τ* is the long-time (i.e. the timescale of hours). Maximum cell velocity obtained during free expansion of the MDCK cell monolayer was 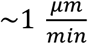 (Serra-Picamal et al., 2012). Long-time oscillatory change of the cell velocity leads to a long-time oscillatory change of strain and corresponding cell residual stress in the form of the propagative waves. These successive extensions and compressions of cell monolayer cause alternate softening and stiffening (Sera-Picamal et al., 2012; Pajic-Lijakobic and Milivojevic, 2020c). The corresponding periodic change of the monolayer rheological behavior is closely connected to the forward backward flows (Serra-Picamal et al., 2012). The stress relaxation occurs at a short-time scale t, i.e. the time scale of minutes (Marmottant et al., 2009; Khalilgharibi et al., 2019). Consequently, stress relaxes during successive short time relaxation cycles under constant strain per cycle (Pajic-Lijakovic and Milivojevic, 2019a;2020b;2020c). The cell displacement field 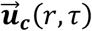 induces generation of shear and volumetric strains. The shear strain is equal to 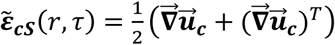, while the volumetric strain is equal to 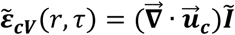. These strains lead to the generation of corresponding shear and normal stresses, their short-time relaxations and the long-time accumulation of cell residual stresses (Pajic-Lijakovic and Milivojevic, 2020b;2021a). The main characteristic of propagative mechanical waves is that (1) cell normal residual stress component *σ*_*cxx*_ and corresponding strain rate 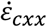 are in phase quadrature, (2) normal stress component is always tensional, and (3) velocity and cell tractions are uncorrelated (Serra-Picamal et al., 2012; Petrolli et al., 2021). These characteristics of the propagative waves pointed to the Zener model suitable for the viscoelastic solid as was discussed by Pajic-Lijakovic and Milivojevic (2020c). The Zener model is expressed as:

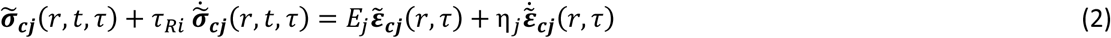

where 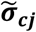 is the shear or normal stress, 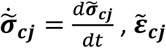 is the shear or volumetric strain, 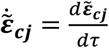 is the strain rate, *E*_*j*_ is the shear or Young’s elastic modulus, and *η*_*i*_ is the shear or volumetric viscosity, and *τ*_*Rj*_ is the corresponding stress relaxation time. Stress relaxation under constant strain 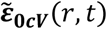 per single short-time relaxation cycle can be expressed starting from the initial 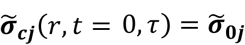 as:

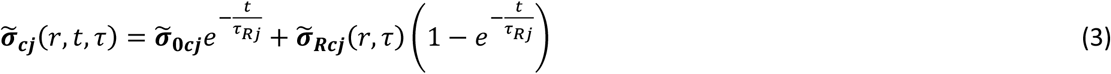

where 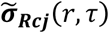 is the residual shear or normal stress equal to 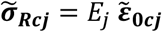. Consequently, the correlation 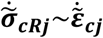 was observed experimentally by Serra-Picamal et al. (2012) which represents a confirmation of the Zener model validity.

The corresponding force-balance responsible for the long-time change of cell velocity has been expressed in the form of inertial wave equation (Pajic-Lijakovic and Milivojevic, 2020c):

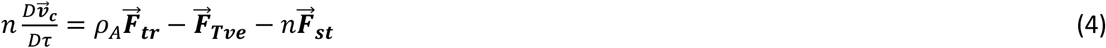

Where 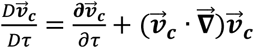 is the material derivative (Bird et al., 1960) and *ρ*_*A*_ is the density of cell-matrix adhesion contacts which depends on the cell type and rheological behavior of matrix. Collision of forward and backward flows can induce change in the cell packing density and consequently change in the state of viscoelasticity (Serra-Picamal et al., 2012; Nnetu et al., 2012).

## 3. Collision between forward and backward flows: the cell packing density increase

Generated forward and backward flows can collide each other during the cell monolayer free expansion. The collision induces an increase in the cell stress within a collision zone (CZ) and a corresponding increase in the cell packing density. The cell packing density increase changes the state of viscoelasticity and can induce the cell jamming state transition (Nnetu et al., 2012; Serra-Picamal et al., 2012). The CZ is labeled as ℛ ± Δ (where Δ is the length order of magnitude larger than the size of a single cell). This zone can move in forward or backward directions depending on the cell velocities 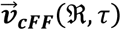 and 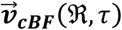 (where 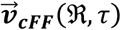 is the cell velocity of a forward flow in the CZ and 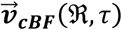 is the cell velocity of a backward flow in the CZ). The resulted cell velocity in the CZ, 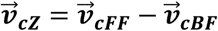 is equal to:

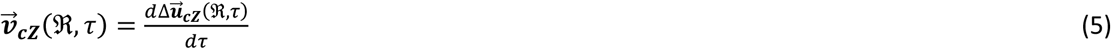

where 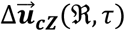 is the resulted cell displacement in the CZ which induces the generation of volumetric strain 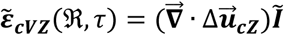 and shear strain 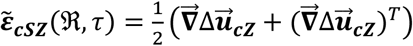. These strains induce generation of corresponding cell normal stress 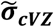 and cell shear stress 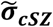. An increase in the cell normal stress leads to an increase in the cell packing density *n*(ℛ, *τ*) (Trepat et al., 2009).

Serra-Picamal et al. (2012) measured the maximum tensional stress in x-direction during the cell monolayer free expansion of ∼300 *Pa*. Tensional stress in one direction induces generation of the compressive stress in the other in order to keep the structural integrity of the cell monolayer parts. An increase in cell compressive residual stress influences the cell state on two ways: (1) directly, by compressing the single cells and (2) indirectly, by increasing the cell packing density and on that base intensifying cell-cell interactions. Single cells well tolerate a compressive stress of several kPa (Kalli and Stylianopoulos, 2018). Recently, Dolega et al. (2021) considered the long-time rearrangement of CT26 mouse cells aggregate within an extracellular matrix under the compression induced by osmotic stress of 5 kPa. They compared the response of cells within two types of experiments, i.e. under global compression and selective compression. Selective compression is induced by the external solution of small dextran molecules while the global compression is induced by the same concentration of large dextran molecules. The selective compression deforms cells while the cell packing density stays approximately constant. The global compression induces single-cell deformation and leads to an increase in the cell packing density. Dolega et al. (2020) revealed that the single-cell compression leads to a decrease in cell volume of ∼ 15 % while cells keep their activity. However, an increase in the cell packing density significantly reduces cell mobility and proliferation. Srivastava et al. (2020) considered response of Dictyostelium cells under compressive stress and pointed out that the compressive stress of 100 Pa is sufficient to cause a rapid (<10 s) change of the way of moving from pseudopods to blebs. Cells are flattened under this compressive stress and lose volume; while the actin cytoskeleton is reorganized such that myosin II molecules are recruited to the cortex. Cells keep their active (contractile) state under this experimental condition. We are interested here in the density driven reduction of cell motility, a phenomenon frequently caused by CCM.

The constitutive stress-strain model for the CZ depends on: (1) the cell packing density, (2) cell mobility, and (3) their inter-relation. An increase in cell density can suppress cell movement and on that base influences the viscoelasticity within a CZ. If the cell movement is totally suppressed, this part of a cellular system undergo to the cell jamming state transition (Nnetu et al., 2013; Pajic-Lijakovic and Milivojevic, 2019c). Pajic-Lijakovic and Milivojevic (2021a,b) proposed several viscoelastic models suitable for various viscoelastic regimes depending on cell packing density and cell velocity based on the experimental data proposed in the literature (Petitjean et al., 2010; Serra-Picamal et al., 2012, Nnetu et al., 2012;2013; Notbohm et al., 2016; Tlili et al., 2018). These findings are systematized and presented in the **Table 1**:

**Table 1.** Various constitutive models correspond to various cell packing density.

| Viscoelastic regimes | $\vec{v}_{cZ}(\mathfrak{R}, \tau) \left( \frac{\mu m}{min} \right)$ | Constitutive model | Volume fraction of passive cells $\phi_r(\mathfrak{R}, \tau)$ | Ordering parameter $\frac{L_c}{L_{cmax}}$ |
| --- | --- | --- | --- | --- |
| 1.Convective Regime | $0.1 \frac{\mu m}{min} < \vec{v}_c$<br>$< \sim 1 \frac{\mu m}{min}$ | The Zener model<br>$\tilde{\sigma}_{cZj}(\mathfrak{R}, t, \tau) + \tau_{Rj} \dot{\tilde{\sigma}}_{cZj} = E_j \tilde{\epsilon}_{cZj} + \eta_j \dot{\tilde{\epsilon}}_{cZj}$<br>Cell residual stress<br>$\tilde{\sigma}_{RcZj} = E_j \tilde{\epsilon}_{cZj}$ | $\sim 0$ | ↓ |
| 2.Conductive regime | $\vec{v}_c \sim 10^{-3} \frac{\mu m}{min}$<br>$- 10^{-2} \frac{\mu m}{min}$ | The Kelvin-Voigt model<br>$\tilde{\sigma}_{cZj}(\mathfrak{R}, \tau) = E_j \tilde{\epsilon}_{cZj} + \eta_j \dot{\tilde{\epsilon}}_{cZj}$<br>The stress cannot relax.<br>$\tilde{\sigma}_{cZj} = \tilde{\sigma}_{RcZj}$ | $\sim 0$ | ↓ |
| 3.Damped conductive regime: Transient sub-regime | $\vec{v}_c \rightarrow 0$ | The Fraction model for the transient state<br>$\tilde{\sigma}_{cZj}(\mathfrak{R}, \tau) = \eta_{\beta(\tau)j} D^{\beta(\tau)}(\tilde{\epsilon}_{cZj})$<br>$\beta(\tau)$ decreases from $\beta = 1/2$ to $\alpha$<br>The stress cannot relax.<br>$\tilde{\sigma}_{cZj} = \tilde{\sigma}_{RcZj}$ | increases from 0 to 1 | ↓ |
| 4.Damped conductive regime: Jamming sub-regime | $\vec{v}_c = 0$ | The Fraction model for the jamming state<br>$\tilde{\sigma}_{cZj}(\mathfrak{R}, \tau) = \eta_{\alpha j} D^{\alpha}(\tilde{\epsilon}_{cZj})$<br>For $0 < \alpha < 1/2$<br>The stress cannot relax.<br>$\tilde{\sigma}_{cZj} = \tilde{\sigma}_{RcZj}$ | $\sim 1$ | ↓ |

where $D^{\alpha} \tilde{\epsilon}(r, \tau) = \frac{d^{\alpha} \tilde{\epsilon}(r, \tau)}{d\tau^{\alpha}}$ and $D^{\beta} \tilde{\epsilon}$ are the fractional derivative, and $\alpha$ is the orders of fractional derivatives, $\eta_{\alpha i}$ is the effective modulus (volumetric or shear) for the transient and jamming sub-regimes. Caputo's definition of the fractional derivative of a function $n(r, \tau)$ was used, and it is given as (Podlubny, 1999): $D^{\alpha} \tilde{\epsilon} = \frac{1}{\Gamma(1-\alpha)} \frac{d}{d\tau} \int_0^{\tau} \tilde{\epsilon}(r, \tau') d\tau'$ (where $\Gamma(1-\alpha)$ is a gamma function).

Maximum cell velocity obtained during free expansion of the MDCK cell monolayer is 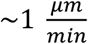 (Serra-Picamal et al., 2012). Nnetu et al. (2012) considered CCM of MCF-10A cell monolayers and obtained the maximum local cell velocity of 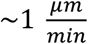 for the low cell packing density of 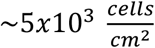 The corresponding viscoelasticity has been described by the Zener model (Pajic-Lijakovic and Milivojevic, 2020c) which is in accordance with the behavior of generated propagative waves, i.e. 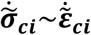 experimentally confirmed by Serra-Picamal et al. (2012). An increase in cell packing density leads to a decrease in cell velocity. Notbohm et al. (2016) considered CCM of the confluent MDCK cell monolayer and pointed to the maximum cell velocity equal to 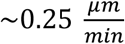. Petitjean et al. (2010) revealed that the MDCK cell monolayer reached the confluence for the cell packing density of 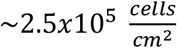 and the cell velocity of 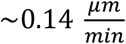. Garcia et al. (2015) considered 2D CCM of various types of cells and emphasized that cell packing density increase from 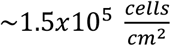 to 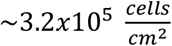 causes a decrease in the cell velocity for an order of magnitude. They revealed that a decrease in cell velocity from 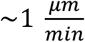 to 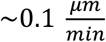 induces a decrease in the velocity correlation length from to and on that base induces the disordering of cell movement.

Pajic-Lijakovic and Milivojevic (2021b) tried to estimate which mechanism of cell movement, convective or conductive, is dominant in this viscoelastic regime. They calculated the convective velocity equal to 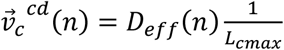 (where *D*_*eff*_(*n*) is the effective diffusion coefficient and *L*_*max*_ is the maximum velocity correlation length). The effective diffusion coefficient decreases from 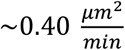 to 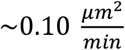 when the packing density of MDCK cells increases from 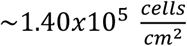 to 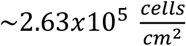 (Angelini et al., 2011). The maximum correlation length for 2D CCM is *L*_*max*_∼150 *µm* (Petroli et al., 2021). Corresponding value of the cell conductive velocity 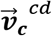 is in the range of 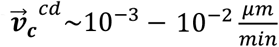 (Pajic-Lijakovic and Milivojevic, 2021b). Consequently, the viscoelasticity described by the Zener model is obtained for the cell velocities 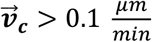 (Nnetu et al., 2012; Serra-Picamal et al., 2012; Notbohm et al., 2016) and corresponds to a convective regime (**Table 1**).

Petrolli et al. (2021) emphasized that the velocity correlation length further decreases when the cell packing density increases. Petitjean et al. (2010) reported that the cell velocity is 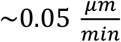 for the packing density of MDCK cells equal to 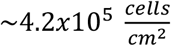, which can correspond to a conductive regime (**Table 1**). Cell movement is significantly suppressed within this regime and cell stress cannot relax. Pajic-Lijakovic and Milivojevic (2021b) proposed the Kelvin-Voigt model for describing the viscoelasticity in the conductive regime.

Further increase in cell packing density additionally reduces cell movement and leads to an anomalous nature of energy dissipation characteristic for the damped-conductive regime (Nnetu et al., 2013; Pajic-Lijakovic and Milivojevic, 2021b). Within this regime, we considered two sub-regimes: (1) the transient sub-regime and (2) the cell jamming sub-regime. Anomalous effects of cell movement in the form of sub-diffusion have been quantified by the damping coefficient equal to *α* ≤1/2 (Pajic-Lijakovic and Milivojevic, 2019c;2021b) (**Table 1**). In the transient sub-regime, the cell velocity tends to zero, i.e.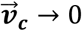 and volume fraction of cells in passive (non-contractile) state increases from 0 to 1. The corresponding velocity correlation length is approximately equal to the single-cell size (Petrolli et al., 2021). Consequently, this sub-regime can be treated as the pseudo-phase transition between active (migrating) cell state to passive (resting) cell state obtained under constant cell packing density (Pajic-Lijakovic and Milivojevic, 2019a; 2020a).

The cell jamming sub-regime also represents a part of the damped conductive regime. Cell rearrangement is additionally damped, while the cell velocity is equal to zero (**Table 1**). Under jamming, all cells are in the passive (non-contractile) state. Bi et al. (2015) treated this cell state transition by the cell shape parameter. Tissues containing cells with a high shape factor (elongated cells) are predicted to be in a motile, unjammed state, while tissues containing cells with a lower shape factor (less elongated) are predicted to be passive, i.e. jammed. Nnetu et al. (2013) pointed to the anomalous nature of energy dissipation during the rearrangement of MCF-10A cell monolayers for the cell packing density of 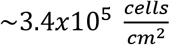. They determined the damping coefficient equal to *α* ∼0.2. This state corresponds to the cell jamming state. Tlili et al. (2018) considered density driven CCM of MDCK cell monolayers. They revealed that cell velocity drops to zero for the cell packing density equal to 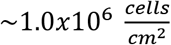.

## 4. Single-cell response under high cell packing density (the damped-conductive regime)

We are interested in cell-cell interactions under very high cell packing density which can induce anomalous effects in a long-time cell rearrangement characteristic for the damped-conductive regime. How single cells reconcile potentially conflicting cues remains poorly understood. Zimmermann et al. (2016) considered the cell jamming state transition during CCM of MDCK cell monolayers. They reported that cells under jamming state actively down-regulate their propulsion forces in response to an increase in cell packing density by favoring the contact inhibition of locomotion (CIL) as the main mechanism. CIL represents cell head-to-head collisions (Mayor and Carmona-Fontaine, 2010). This type of collisions can suppress forward locomotion on cell–cell contact and (in many but not all cell types that display it) is followed by protrusion collapse and a subsequent redirection of cell motility (Roycroft and Mayor, 2016). Interestingly, many malignant cells do not display CIL when interacting with other cell types (heterotypic), but retain CIL in interactions within a same cell type (homotypic) (Zimmermann et al., 2016). Several effects have been recognized within cellular system near jamming: (1) a decrease in the propulsion force produced by each cell, (2) an increase in the friction of the cells with the substrate, and (3) increase in the cell−cell effective friction (Garcia et al., 2015).

In order to analyze cell-cell interactions, we introduced the dimensionless criteria 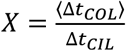 (where ⟨Δ*t*_*COL*_⟩ is the average time between two cell collisions, Δ*t*_*CIL*_ is the characteristic time for single-cell repolarization equal to Δ*t*_*CIL*_ ≈ 10 *min* (Alert et al., 2019). The average time between two cell collisions depends on: (1) the cell packing density *n*(ℛ, *τ*), (2) ordering trend of cell movement expressed as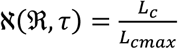 (**Table 1**) (where *L*_*C*_ is the velocity correlation length and *L*_*max*_ is the maximum cell correlation length), and (3) the average cell speed 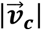 expressed in the form of effective temperature by Pajic-Lijakovic and Milivojevic (2019c;2021a). They introduced the relationship 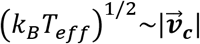 (where *T*_*eff*_ is the effective temperature and *k*_*B*_ is the Boltzmann constant). Consequently, the change of the dimensionless criteria, responsible for changing the mechanical state of single-cells, can be expressed as:

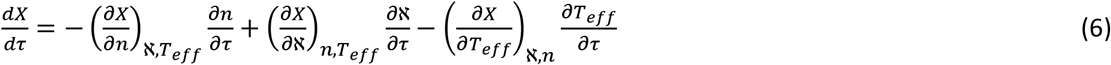

An increase in cell packing density under constant effective temperature and the system ordering parameter ℵ leads to a decrease in the time ⟨Δ*t*_*COL*_⟩. A decrease in cell ordering parameter ℵ under constant cell packing density and effective temperature *T*_*eff*_ induces an increase in cell-cell collisions and on that base a decrease in the average collision time. An increase in the cell mobility expressed by the effective temperature under constant cell packing density and the system ordering parameter leads to a decrease in the time ⟨Δ*t*_*COL*_⟩.

If the dimensionless criteria *X* is *X* ≥ 1 cells have enough time to re-polarize and adapt under micro-environmental conditions. However, the dimensionless criteria *X* becomes *X* < 1 for the cells near jamming. Consequently, cells are unable to re-polarize and stay frozen in the transition state. If this state lasts longer time period (the time scale of hours), cells reduce the activity of stress fibers (Zimmermann et al., 2016) which leads to the cell pseudo-phase transition from active to passive cell state. Pajic-Lijakovic and Milivojevic (2021b) defined the cell packing density under jamming state as: 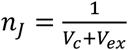 (where *V*_*C*_ is the average single-cell volume and *V*_*ex*_ is the excluded volume per single cell). The excluded volume is expressed based on the second virial coefficient 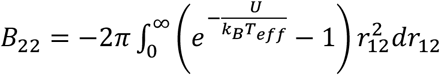 (where *U*(*r*_12_) is the interaction potential between two cells). They pointed out that when the potential *U*(*r*_12_) → ∞, the second virial coefficient tends to the cell excluded volume, i.e. *B*_22_ → *V*_*ex*_ The condition *X* < 1 is reached in the damped-conductive regime and leads to the cell pseudo-phase transition described by an increase in the volume fraction of passive (resting) cells within a CZ which will be discussed based on thermodynamic and rheological approaches.

### 4.1 The pseudo-phase transition near cell jamming

The volume fraction of passive cells ϕ_*r*_ (ℛ, *τ*) within a CZ is ϕ_*r*_ (ℛ, *τ*) → 0 for convective and conductive regimes (**Table 1**). However, damped cell rearrangement achieved in the transient sub-regime leads to an increase in the volume fraction of cells in the passive (resting) state from 0 to 1. This sub-regime can be treated as a pseudo-phase transition from cell active (contractile)-to passive (non-contractile) state. Corresponding increase in the ϕ_*r*_ (ℛ, *τ*) for the transient sub-regime can be expressed as:

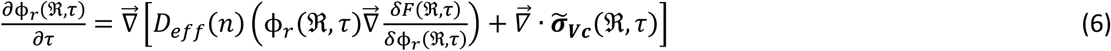

where *D*_*eff*_ (*n*) is the effective diffusion coefficient, *n*(ℛ, *τ*) is the cell packing density, *F* (ℛ, *τ*) is the free energy functional which accounts for the action of dynamic surface tension (where, *F* (ℛ, *τ*) = ∫ *γ*, (ℛ, *τ*) *d*^*2*^*r* and *γ*(ℛ, *τ*) is the surface tension). The first term on the right hand side represents the thermodynamic affinity which accounts for cumulative effects of biochemical processes such as cell adhesion, signalling and gene expressions, while an increase in the gradient of cell normal residual stress (i.e. the second term) can suppress cell movement which leads to an increase in the volume fraction of passive cells. The volume fraction ϕ_*r*_ (ℛ, *τ*) becomes equal to ϕ_*r*_ (ℛ, *τ*) = 1 and stays constant within the cell jamming sub-regime.

## 5. The viscoelasticity regimes and the stiffness of the collision zone

An increase in the volume fraction of passive cells leads to the CZ softening for the transient sub-regime. Active (contractile) cells are stiffer than passive (non-contractile) ones due to an accumulation of the contractile energy (Kollmannsberger et al., 2011; Pajic-Lijakovic and Milivojevic, 2019a;2020b). Schulze et al. (2017) reported that Young’s modulus of contractile MDSC cell monolayer ∼33.0 ± 3.0 *kPa* is, while the modulus of non-contractile cells is significantly lower and equal to ∼15.6 ± 5.5 *kPa*. Lange and Fabry (2013) reported that muscle cells can change their elastic modulus by over one order of magnitude from less than 10 kPa in a relaxed (resting) state to around 200 kPa in a fully activated (contractile) state. Consequently, the single-cell stiffness has been quantified by various modulus, such as: (1) complex shear modulus 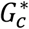 (Trepat et al., 2004), (2) complex longitudinal modulus 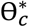 (Ferna’ndez et al., 2006), and (3) differential modulus 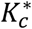 (Kollmannsberger et al., 2011) under various externally applied stress or strain conditions. These contributions pointed to the relationship between cell pre-stress and the single-cell stiffness in the form |*M*_*C*_ |∼ (*σ*_*p*_ + *σ*_*a*_)^*β*^ (where | *M*_*C*_ | is the argument of complex modulus equal to 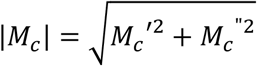, is the storage modulus of single-cell, *M*_*c*_” is the loss modulus of single cell, *σ*_*a*_ is the stress of passive cell, *σ*_*p*_” is the cell active contractile pre-stress, and *β* is the exponent). Notbohm et al. (2016) correlated cell pre-stress with the concentration of phosphorylated myosin.

The stiffness of multicellular systems can be quantified by the argument of the complex modulus (shear and volumetric) 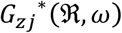, i.e. 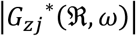. The complex modulus 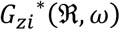 can be expressed from the constitutive models presented in **Table 1** for various viscoelastic regimes obtained by Fourier transform 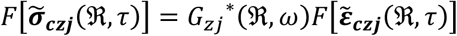. The shear or volumetric complex modulus *G*_*zj*_^*^ is equal to *G*_*zj*_^*^ = *G*_*zj*_^′^ (where *G*_*zj*_^”^ is the storage modulus, *ω* is the loss modulus, is the angular velocity, and 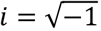). The argument of the complex modulus for various viscoelastic regimes is shown in **Table 2**.

**Table 2.** The argument of the complex modulus for various viscoelastic regimes as a measure of the system stiffness

| Viscoelastic regimes | The argument of the complex modulus $\vert G_{zj}^*(\Re, \omega)\vert $ |
| --- | --- |
| (1) Convective regime | $\frac{\sqrt{(E_j + \eta_j \tau_{Rj} \omega^2)^2 + (\eta_j \omega - E_j \tau_{Rj} \omega)^2}}{1 + \tau_{Rj}^2 \omega^2}$ |
| (2) Conductive regime | $\sqrt{E_j^2 + \eta_j^2 \omega^2}$ |
| (3) Damped conductive regime: Transient sub-regime | $\eta_{1/2j} \omega^{1/2} \geq \vert G_{zj}^*(\Re, \omega)\vert _T > \eta_{\alpha j} \omega^\alpha$ |
| (4) Damped conductive regime: Jamming sub-regime | $\eta_{\alpha j} \omega^\alpha$ |

If the collision between forward and backward flows is capable of inducing the damping-conductive regime, the cells undergo active to passive cell state transition at the line of contact ℛ. Under this condition, the volume fraction of cells in the passive state increases frontally to both sides ℛ ± Δ. The transient regime corresponds to a so called “sandwich structure” presented schematically in Figure 2.

**Figure 2.**
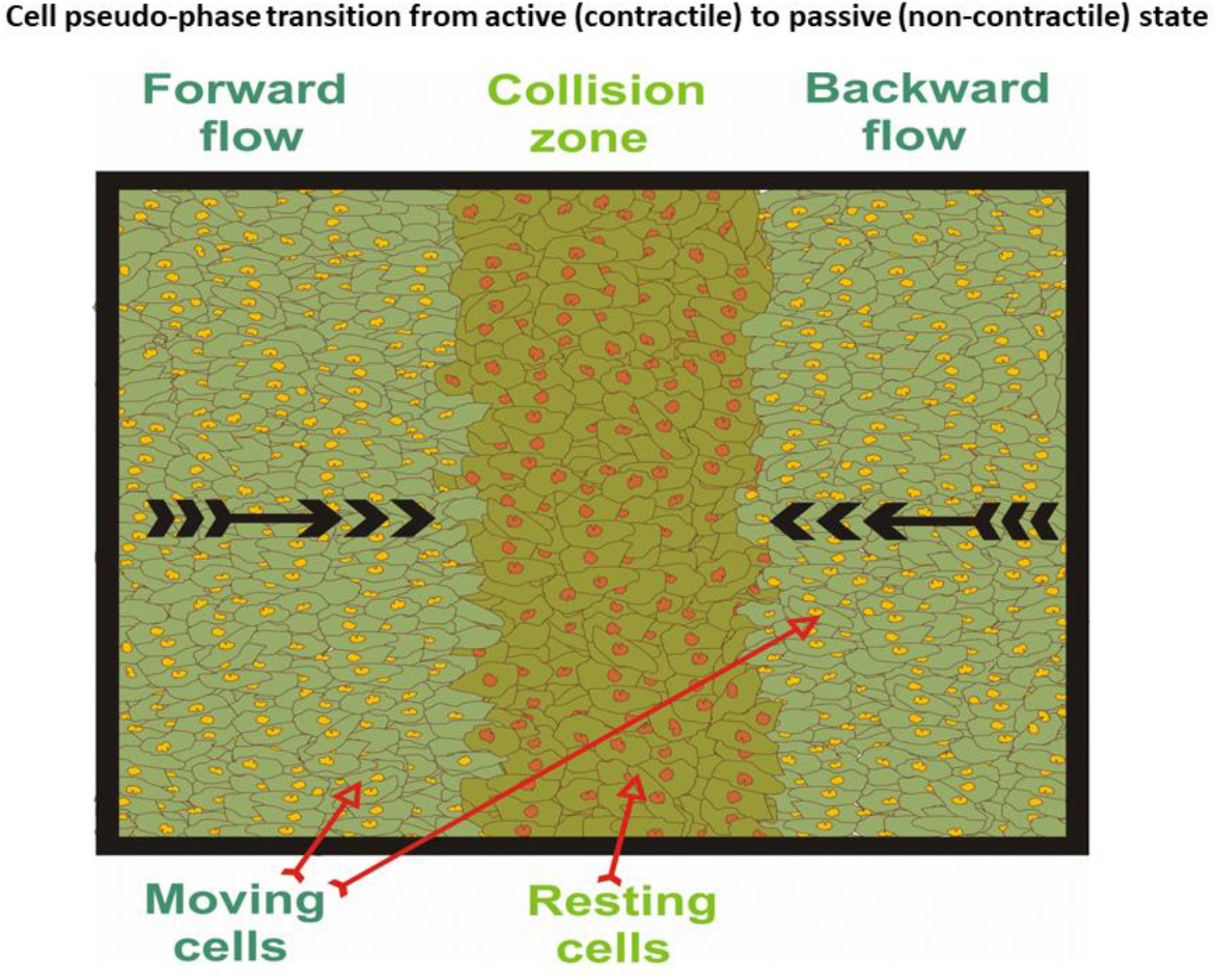
Schematic presentation of an increase in volume fraction of cells in the resting state within a collision zone.

Cells in the passive state are placed in the internal region, while the active cells are placed in the boundary regions of the collision zone. Consequently, the viscoelasticity of the CZ depends on (1) the volume fraction of migrating cells, ϕ_*m*_, (2) the volume fraction of passive (resting) cells ϕ_*r*_ such that ϕ_*r*_ = 1 − ϕ_*m*_, (3) the viscoelasticity of migrating cells 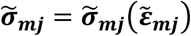 (where 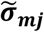 is the shear or normal stress generated within a migrating cell pseudo phase and 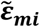 is the corresponding shear and volumetric strain), and (4) the viscoelasticity of passive (resting) cells 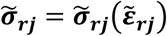 (where 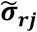 is the shear or normal stress generated within a resting cell pseudo-phase and 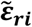 is the corresponding shear and volumetric strain), (Pajic-Lijakovic and Milivojevic, 2019a;2020a). This “sandwich structure” of cell-pseudo-phases pointed to parallel mode coupling described by Pajic-Lijakovic and Milivojevic (2019a;2020a). The resulted constitutive model for the transient sub-regime for the parallel mode coupling between migrating and resting cell pseudo-phases can be expressed as (Pajic-Lijakovic and Milivojevic, 2019a,2020a):

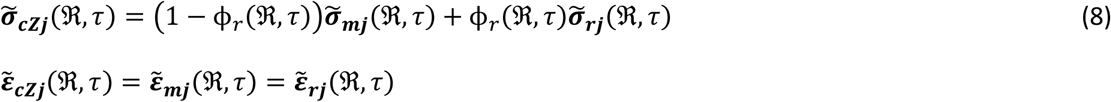

where 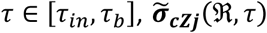 is the shear or normal stress within the CZ expressed by the fractional constitutive model presented in **Table 1** and 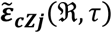 is the corresponding shear or volumetric strain. The initial condition for the transient sub-regime is: at time *τ*_*in*_, the volume fraction of passive (resting) cells in the CZ is ϕ_*r*_ (ℛ, *τ*_*in*_) = 0 and the total stress is equal to 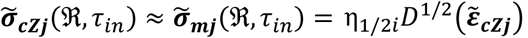. The volume fraction of resting cells increase with time under constant cell packing density and at the boundary time *τ*_*b*_ = *τ*_*in*_ + Δ*τ* is equal to ϕ_*r*_ (ℛ, *τ*_*b*_) → 1. This condition corresponds to a cell jamming state. The total stress at *τ*_*b*_ = *τ*_*in*_ is equal to 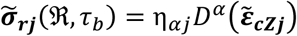 for 0 < *α* 1/2.

The stiffness of the collision zone |*G*_*zj*_^*^ | depends on: (1) the cell mobility quantified by the effective temperature, (2) the cell packing density, (3) the volume fraction of cells in the resting state, and (4) the single-cell stiffness. The effective temperature decreases and the cell packing density increases from convective to conductive regime, while single-cells keep their active (contractile) state. These changes lead to an increase in the CZ stiffness as reported by Schierbaum et al (2017). However, the volume fraction of resting cells increase within the transient sub-regime from 0 to 1 and single-cells change their state from contractile to non-contractile. This change of the viscoelasticity from conductive regime to the transient sub-regime leads to the CZ softening. Under jamming state, all cells within the CZ are in the passive state. Consequently, the maximum stiffness is achieved in the conductive regime, while the minimum stiffness corresponds to the cell jamming sub-regime. The stiffness change within a CZ is expressed in the dimensionless form 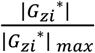 and presented schematically in Figure 3.

**Figure 3.**
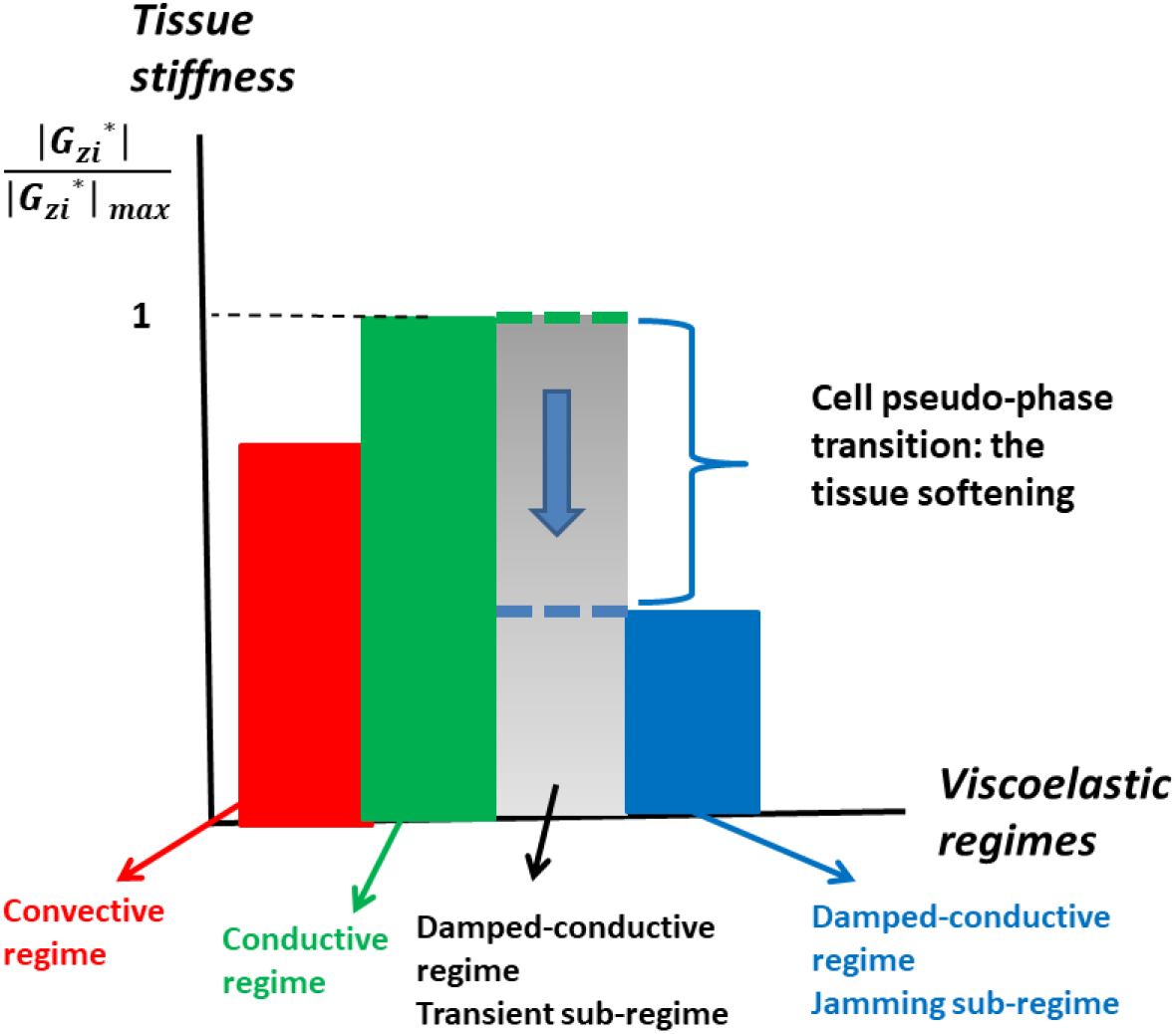
Change the stiffness within a collision zone as a consequence of an increase the cell packing density from the convective regime to the cell jamming regime.

The stiffness increases from convective to conductive regime, reaches the maximum value in the conductive regime and then decrease through the transient sub-regime as the consequence of the cell pseudo-phase transition from active (contractile) to passive (non-contractile) caused by high cell packing density. The stiffness reaches the minimum value under the cell jamming state. Cell unjamming state transition is related to an increase in the volume fraction of active (contractile) cells in the boundary region of the CZ after a time period of hours.

## 6. Conclusion

Significant attempts have been made to discuss the density-driven change in a long-time cell rearrangement. Less attention has been paid clarify the role of viscoelasticity caused by CCM in this process. Despite extensive research devoted to the study of the long-time cell rearrangement under in vitro and in vivo conditions we still do not understand the process from the standpoint of rheology. An increase in the cell packing density changes the tissue viscoelasticity and induces an increase in the tissue stiffness if and only if cells keep their active (contractile) mechanical state. However, cells near jamming undergo the pseudo-phase transition from active (contractile) to passive (non-contractile) state. Active cells are much stiffer than passive ones due to an accumulation of the contractile energy. Consequently, the cell pseudo-phase transition has to be included into the corresponding constitutive model.

The phenomenon is discussed on the experimentally well elaborated model system such as the cell monolayer free expansion. The monolayer free expansion induces the generation of mechanical waves. Mechanical waves represent oscillations of cell velocity and the relevant rheological parameters. The cell velocity oscillations in the form of forward and backward flows are driven by the viscoelastic force, surface tension force, and traction force. Collision of the forward and backward flows induces the cell packing density increase as well as the change in the state of viscoelasticity within a CZ which can lead to a cell jamming state transition. Various viscoelastic regimes are discussed such as: (1) convective regime based on the Zener model (for the cell velocity in the range of 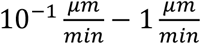), conductive regime based on the Kelvin-Voigt model (for the cell velocity in the range of 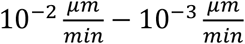), and (3) damped conductive regime based on the Fractional model. The damped-conductive regime accounts for the transient sub-regime (for the cell velocity tends to zero) and the jamming sub-regime (for the cell velocity equal to zero). The transient sub-regime is the prerequisite for cells to achieve the jamming state. This sub-regime accounts for the cell pseudo phase transition from active (contractile) to passive (non-contractile) state. This change of the viscoelasticity from regime to regime was discussed from the standpoint of the tissue stiffness expressed based on proposed viscoelastic models.

The main conclusion was extracted, based on modeling consideration, that the tissue stiffness increases from convective to conductive regime, reaches the maximum value in the conductive regime and then decrease through the transient sub-regime as the consequence of the cell pseudo-phase transition from active (contractile) to passive (non-contractile). The stiffness reaches the minimum value under the cell jamming state.

## Acknowledgement

This work was supported by the Ministry of Education, Science and Technological Development of the Republic of Serbia (Contract No. 451-03-9/2021-14/200135).

